# *Arabidopsis thaliana* RHAMNOSE 1 condensate formation drives UDP-rhamnose synthesis

**DOI:** 10.1101/2024.02.15.580454

**Authors:** Sterling Field, Yanniv Dorone, Will P. Dwyer, Jack A. Cox, Renee Hastings, Madison Blea, Olivia M. S. Carmo, Dan Raba, John Froehlich, Ian S. Wallace, Steven Boeynaems, Seung Y. Rhee

## Abstract

Rhamnose is an essential component of the plant cell wall and is synthesized from uridine diphosphate (UDP)-glucose by the RHAMNOSE1 (RHM1) enzyme. RHM1 localizes to biomolecular condensates in plants, but their identity, formation, and function remain elusive. Combining live imaging, genetics, and biochemical approaches in *Arabidopsis* and heterologous systems, we show that RHM1 alone is sufficient to form enzymatically active condensates, which we name ‘rhamnosomes’. Rhamnosome formation is required for UDP-rhamnose synthesis and organ development. Overall, our study demonstrates a novel role for biomolecular condensation in metabolism and organismal development, and provides further support for how organisms have harnessed this biophysical process to regulate small molecule metabolism.

**One-Sentence Summary:** Condensation of RHM1 drives UDP-rhamnose synthesis during plant development.

## Introduction

Rhamnose is an essential building block of the plant cell wall (*1, 2*). It is synthesized in the cytoplasm as the sugar nucleotide uridine diphosphate (UDP)-rhamnose from UDP-glucose by one of three biochemically redundant cytoplasmic enzymes, RHAMNOSE 1, 2, 3 (RHM1, RHM2, RHM3) (*3*). All three enzymes are composed of two catalytically active domains: the N-terminal dehydration domain (Binding Domain 1; BD1) and a C-terminal epimerization-reduction domain (Binding Domain 2; BD2) (*4*), connected by an intrinsically disordered region. The main difference between these three RHMs is their expression across tissues and developmental stages (*5*). RHM1 is the most abundant and ubiquitously expressed RHM enzyme in *Arabidopsis* (*5*). *rhm1* mutants have morphological phenotypes such as twisted petals and roots (*6*). RHM2 is primarily expressed in developing seeds and *rhm2* mutants are associated with defects in seed mucilage synthesis (*4, 7*). RHM3 is poorly characterized and does not have any reported phenotypes.

RHM1 activity is thought to be transcriptionally regulated, where tissues that require more rhamnose synthesis increase expression of *RHM1*. Besides transcriptional control, protein activity could also be regulated through changes in its subcellular localization (*8–11*).

Intriguingly, RHM1 localizes to cytoplasmic bodies (*12, 13*), but their exact identity and potential function, or lack thereof, remains controversial. One study proposed that RHM1 localized to cytosolic stress granules upon heat shock (*12*) when the plant does not need to synthesize rhamnose and the enzyme is likely inactive. Another study found that RHM1 formed foci in dividing petal cells (*6*) when cells need to increase rhamnose production. These two observations suggest RHM1 condensation could potentially affect its enzymatic activity positively or negatively, but evidence for this remains lacking.

Here, we set out to resolve this outstanding question and test directly whether RHM1 condensation has functional importance to plant development. We employed cell biological and analytical chemistry approaches in heterologous systems to show that RHM1 is sufficient for condensate formation and that its condensation is necessary for UDP-rhamnose synthesis. Using genetic approaches, we show that condensate formation is required for cell wall synthesis and organ development *in planta*. In all, we identified a novel biomolecular condensate that we call the ‘rhamnosome’, which is required for the biocatalysis of a critical cell wall building block and is essential for plant development.

### RHM1-bodies are a novel biomolecular condensate

Previous work suggested that RHM1 is a potential stress granule (SG) protein (*12*). To corroborate this, we transiently co-expressed RHM1-GFP in tobacco epidermal cells with the SG marker UBP1c-RFP (*14*). The core SG marker UBP1c robustly formed granules under heat, UV, and oxidative stress (Fig. S1A). In contrast to UBP1c, RHM1 only formed bodies in response to heat stress (Fig. 1A) and RHM1 remained diffuse during UV or oxidative stress (Fig. S1A). Further, the RHM1-bodies that formed during heat stress were clearly distinct from SGs (Fig. 1A). RHM1-bodies were occasionally found next to SGs, suggesting these different condensates may superficially interact, as has been observed for other condensates (*15, 16*). This could explain why RHM1 was previously misidentified as a component of SGs in immunoprecipitation assays (*12*). Since RHM1 localized to a distinct and uncharacterized condensate *in planta*, we were curious if RHM1 could be a scaffold protein sufficient for condensate formation. To test this, we expressed RHM1 in heterologous systems that lacked a rhamnose biosynthesis pathway. RHM1 expressed in yeast and in human cells spontaneously formed RHM1-bodies under non-stress conditions (Fig. 1B and C), indicating that RHM1 does not require cofactors specific to rhamnose-synthesizing organisms to form condensates. We additionally tested if RHM1 localized to SGs in human cells or if RHM1 remained a distinct body as it does *in planta*. Like our tobacco experiments, RHM1 did not localize to SGs in human cells (Fig. 1D), and RHM1-bodies were distinct from SGs (Fig. S1B and C). We conclude from these findings that RHM1-bodies are not SGs but a novel condensate.

**Figure 1.**
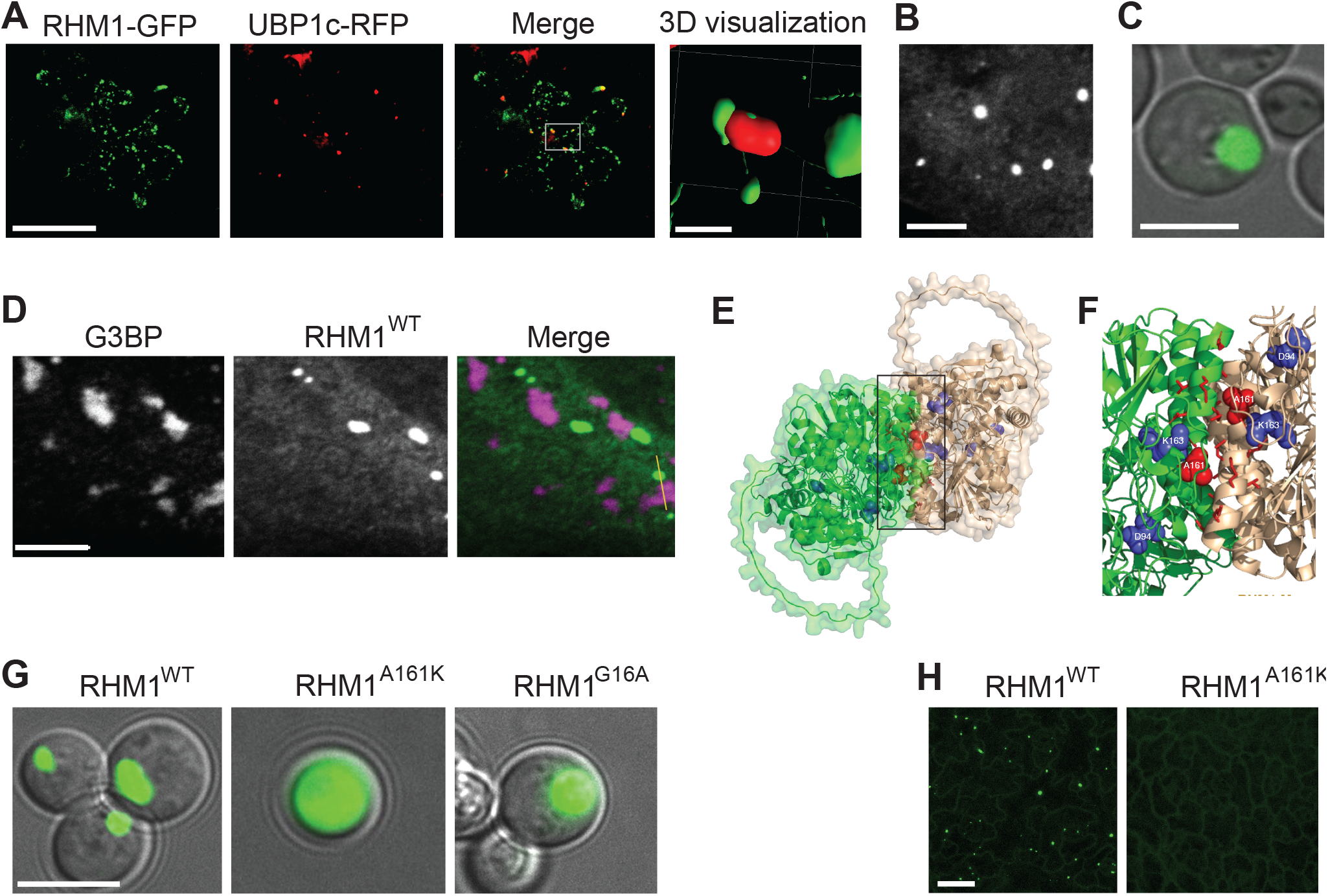
RHM1-bodies are distinct cellular condensates which require RHM1 as a scaffold protein. **A**) Transient expression of RHM1-GFP with the stress granule marker UBP1c-RFP in *Nicotiana benthamiana* after 1 hr of 42°C heat stress. Each image is a maximum projection of 20 images captured in the Z-direction to visualize a whole cell. *Right*, 3D visualization of rhamnosomes and a UBP1c-marked stress granule. Scale bars are 40 μm (*left*) and 3 μm (*right*). **B**) RHM1 transiently expressed in unstressed human U2OS cells (scale bar = 5 μm). **C**) RHM1 expressed in unstressed *Saccharomyces cerevisiae* W303a cells (scale bar = 5 μm). **D**) RHM1 transiently expressed in U2OS cells treated with sodium arsenite and probed with the immunofluorescence stress granule marker anti-G3BP. Yellow line indicates pixels used for linescan (see Fig. S1B; scale bar = 5 μm). **E** and **F**) *In silico* prediction of RHM1 dimerization using a space filling model. *E*, dimerization of two RHM1 monomers (monomers are shown as green and tan). Boxed region is magnified in F. *F*, the dimerization interface between two RHM1 monomers. Red residues are at the interaction surface, with the A161 residue highlighted by space filling. Blue space filled residues are those previously identified to be critical for RHM1 catalytic function (*4*). **G**) Expression of RHM1-GFP and RHM1 point mutations in unstressed yeast cells (scale bar = 5 μm). **H**) Expression of *pRHM1*::RHM1-GFP and *pRHM1*::RHM1^A161K^-GFP in unstressed developing leaves of stable *Arabidopsis* transgenic lines (scale bar = 25 μm). All micrographs in this figure are representative of at least 3 independent experiments.

### RHM1-body formation requires the RHM1 homodimerization domain

Since RHM1 is sufficient to spontaneously form RHM1-bodies in a heterologous system, we next asked which protein domains of RHM1 were essential for RHM1 condensation. RHM1 forms homodimers through a pair of alpha helices in BD1 (*17*), which have hydrophobic residues exposed at the interaction surface (*17*). To test if RHM1 condensation requires dimerization through this hydrophobic interaction mediated by the pair of alpha helices in BD1, we used *in silico* modeling of RHM1 and *in vivo* expression of a RHM1 point mutant. Alanine 161 is at the dimerization surface (Fig. 1E and F), which we mutated to lysine (RHM1^A161K^; Fig. S2A). The RHM1^A161K^ point mutant was not predicted to impact the protein structure of RHM1 (Fig. S2B and C). When expressed in yeast, RHM1^A161K^ was unable to form condensates (Fig. 1G, Fig. S2D). Thus, a surface-exposed hydrophobic patch seems essential for spontaneous RHM1 condensation in our heterologous system, and provides us with a genetic tool to control RHM1 condensation in cells.

To dissect the role of RHM1 condensation *in vivo*, we next asked whether we can prevent RHM1 condensation in *Arabidopsis*. We generated stable transgenic *Arabidopsis* lines expressing either GFP-tagged RHM1 or RHM1^A161K^ under control of the native RHM1 promoter (RHM1-GFP and RHM1^A161K^-GFP) in the loss-of-function *rhm1-2* background. RHM1-GFP formed condensates in developing tissues while RHM1^A161K^-GFP did not (Fig. 1H) despite both proteins being expressed to similar levels (Fig. S3A). These results indicate that the same interaction surface we identified in the heterologous system drives condensation in the physiological context, allowing us to test for the functional consequences of RHM1 condensation *in vivo*.

### RHM1 condensates are tuned by the developmental stage and required for proper tissue morphogenesis

To understand the role of RHM1 condensation in *Arabidopsis*, we first asked where and when RHM1 condensation occurs during development. Prior literature suggested that RHM1 forms puncta in petal cells (*6*). Since petal cells are well-known to undergo extensive cell expansion, and thus require increased cell wall synthesis, we wondered if RHM1 condensation would follow developmental patterns of cell expansion. To characterize the temporal and spatial dynamics of RHM1 condensation during petal development, we examined subcellular and tissue-level localization of the RHM1-GFP protein in petals using live imaging (Fig. 2A). Early in petal development (stage 11, (*18*)), RHM1-GFP had a diffuse signal in the cytoplasm. As the petal expanded (stage 12-13, (*18*)), RHM1-GFP formed bodies in most petal epidermal cells (∼85 % of the cells; Fig. 2A and B). The fraction of cells with RHM1-bodies decreased as petal cells matured (to ∼25 %, stage 14 (*18*)) and senesced (∼3 % of the cells; Fig. 2B). In contrast to RHM1-GFP that formed condensates during petal development, RHM1^A161K^-GFP did not form bodies in petals (Fig. S3B). Together, these data indicate that RHM1 condensation is developmentally regulated in petal cells and correlates with their expansion.

**Figure 2.**
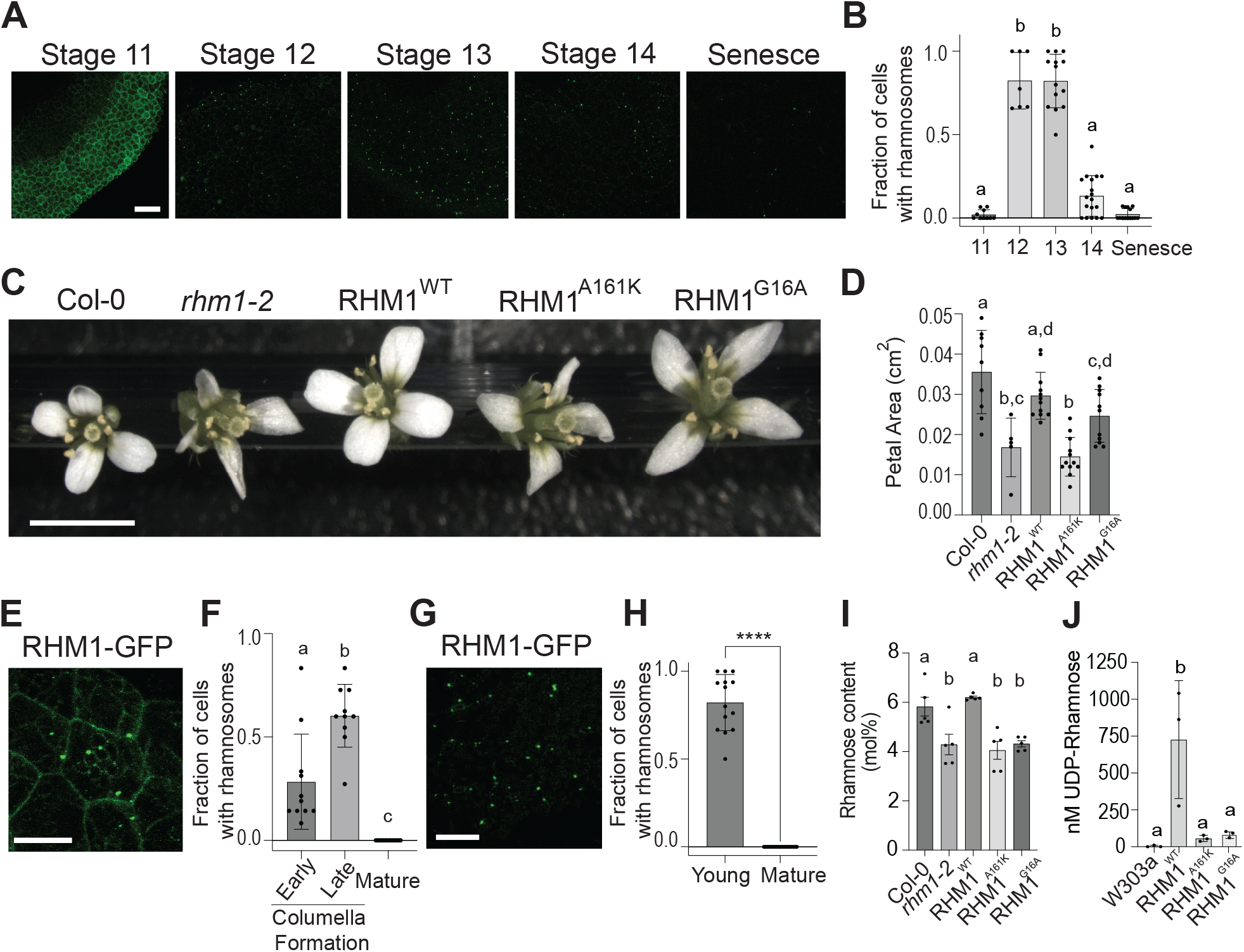
Rhamnosome presence is associated with expanding cells of various organs and is essential for proper plant development. **A**) Confocal microscopy on petals of *Arabidopsis* plants expressing p*RHM1*::RHM1-GFP. Stages of petal development are listed above each image. All images are the same magnification (scale bar = 25 μm). **B**) Quantification of the fraction of petal cells with rhamnosomes during petal development in a 40 μm x 40 μm section of petal tissue. Data is from 7-18 petals, from 3 independent experiments. For statistical significance, different letters represent significant differences (p-value < 0.05, ANOVA). **C**) Comparison of flowers of Col-0, *rhm1-2*, and three *RHM1* promoter driven constructs in the *rhm1-2* background: p*RHM1*::RHM1-GFP, p*RHM1*::RHM1^A161K^-GFP, and p*RHM1*::RHM1^G16A^-GFP (scale bar = 5 mm). **D**) Quantification of petal tissue area from Col-0, *rhm1-2*, and three independent transgenic lines of each construct from C. For statistical significance, different letters represent significant differences (p-value < 0.05, ANOVA). *E*) Rhamnosome formation in the developing seed coat based on p*RHM1*:RHM1-GFP expression (scale bar = 25 μm). **F**) Quantification of the fraction of seed coat cells with rhamnosomes during seed coat development. Data is from 10-14 seeds, from 3 independent experiments. For statistical significance, different letters represent significant differences (p-value < 0.05, ANOVA). **G**) Rhamnosome formation in young leaves based on p*RHM1*::RHM1-GFP expression (scale bar = 25 μm). **H**) Quantification of the fraction of cells with rhamnosomes in leaves. Data is from 16-17 leaves, from 3 independent experiments. Asterisk indicates statistical significance (p-value < 0.05, t-test). **I**) Quantification of the percentage of rhamnose in the cell wall in seedlings of *rhm1-2*, p*RHM1*::RHM1-GFP, p*RHM1*::RHM1^A161K^-GFP, and p*RHM1*::RHM1^G16A^-GFP. Data is from 5 biological replicates. For statistical significance, different letters represent significant differences (p-value < 0.05, ANOVA). **J**) Quantification of 3 independent experiments. UDP-rhamnose synthesized from the yeast lines in Fig 1G after 48 hr of expression. Data is from 3 independent experiments. For statistical significance, different letters represent significant differences (p-value < 0.05, ANOVA). All micrographs in this figure are representative of data from 3 independent experiments.

We next tested if RHM1 condensation was required for petal development at the cellular and organ levels by leveraging the RHM1^A161K^ -GFP line, which lacks RHM1-bodies. *rhm1* mutants have defects in petal development both at the organ and cellular levels. *rhm1* petals are reduced in size and display a helical twist (*6*). At the cellular level, petal epidermal cells (termed cone cells) in *rhm1* develop nanoridges in a parallel pattern, instead of a radial pattern observed in Col-0 wild type plants (*6*). We observed that complementation with RHM1-GFP rescued the *rhm1-2* petal and cone cell phenotypes (Fig. 2C and D, Fig. S3C and D). Since RHM1-GFP is sufficient to rescue the *rhm1-2* phenotypes, we next asked whether the RHM1^A161K^-GFP mutant could rescue the *rhm1-2* petal and cone cell phenotypes. RHM1^A161K^-GFP failed to rescue both phenotypes of *rhm1-2* flowers (Fig. 2C and D, Fig. S3C and D) despite the RHM1^A161K^-GFP protein being expressed to a similar level as RHM1-GFP (Fig. S3A). We reasoned that there were two explanations for why the RHM1-body null construct would not complement *rhm1-2*: either the RHM1-body null protein was unable to synthesize UDP-rhamnose, or RHM1-bodies were required for other processes in petal cells unrelated to UDP-rhamnose synthesis. To test if RHM1-bodies that lacked enzymatic activity could complement the petal phenotype, we generated a catalytically dead RHM1 mutant (RHM1^G16A^-GFP) based on previously identified mutants in rhamnose synthesis (*4*). Glycine 16 is in the BD1 catalytic site, which resides in an internal pocket and is not likely to be involved in dimerization. As expected, RHM1^G16A^-GFP was able to form RHM1-bodies (Fig. 1G and Fig. S2D). RHM1^G16A^-GFP partially rescued the petal phenotype of *rhm1-2* in multiple lines (Fig. 2C and D, Fig. S3C), though it did not rescue the *rhm1-2* cone cell phenotype (Fig. S3D). Together, we conclude that RHM1-bodies are necessary and partially sufficient for petal development and necessary but not sufficient for cone cell wall patterning.

Genetic evidence that RHM1-bodies are required for petal development and cone cell wall patterning prompted us to ask if RHM1-bodies are critical for synthesizing cell wall polymers in general. *Arabidopsis* primary cell walls and seed coat mucilage contain ∼20 % and ∼40 % rhamnose-containing polysaccharides, respectively, primarily in the form of pectic rhamnogalacturonan-I (RG-I) (*19, 20*). We first tested whether RHM1 forms condensates in the developing seed, when seed coat mucilage is synthesized. We observed that RHM1-bodies were present in seed coat cells (Fig. 2E and Fig. S4A) during cellular stages associated with mucilage synthesis (Fig. 2F), and RHM1-bodies were distinctly visible from seed autofluorescence (Fig. S4A). To assay for altered mucilage synthesis in an *rhm1* mutant, we measured the thickness of the *rhm1-2* seed mucilage according to established protocols (*21*). The inner adhesive mucilage layer in *rhm1-2* seeds was substantially reduced compared to Col-0 wild type seeds (Fig. S4B and C). The reduced mucilage in *rhm1-2* was rescued by expressing RHM1-GFP but both the RHM1-body null (RHM1^A161K^-GFP) and the catalytically dead (RHM1^G16A^-GFP) mutant lines failed to rescue this phenotype (Fig. S4B and C). Next, we asked if RHM1-bodies formed during leaf cell growth, which requires cell wall synthesis. As expected, RHM1-bodies were abundant in expanding leaves (Fig. 2G and H). In line with the petal and seed mucilage phenotypes, *rhm1-2* had a reduced total leaf area compared to Col-0, which was rescued by RHM1-GFP but not by the RHM1-body null mutant protein (Fig. S4D and E). Because we observed a reduction in seed mucilage and leaf growth, we next compared the amount of cell wall-associated rhamnose via neutral monosaccharide analysis in *rhm1-2* plants, and *rhm1-2* plants expressing RHM1, RHM1^G16A^, and RHM1^A161K^. *rhm1-2* plants expressing RHM1-GFP had drastically more cell wall-associated rhamnose than *rhm1-2* plants expressing RHM1^A161K^ or RHM1^G16A^ lines, both of which had ∼1/3 less than RHM1-GFP and were indistinguishable from *rhm1-2* (Fig. 2I). In all, RHM1-bodies are readily observed under physiological conditions *in vivo* and specifically in cells and tissues during developmental stages that require extensive cell wall and UDP-rhamnose synthesis. Moreover, a genetic mutant that interferes with RHM1 condensate formation phenocopies an established catalytically-dead mutant, suggesting that RHM1 condensation contributes to RHM1 activity required for proper cell morphology and organ growth via the synthesis of mucilage and rhamnose-containing cell wall polymers.

### RHM1-body disruption perturbs UDP-rhamnose synthesis

Since RHM1-bodies seem required for proper cell and organ development, we next asked whether RHM1 condensation was linked to active UDP-rhamnose synthesis. To answer this question, we chose *Saccharomyces cerevisiae* as a heterologous system to assay UDP-rhamnose synthesis. Yeast does not make rhamnose and allows us to very precisely measure the enzymatic output of this plant biosynthetic pathway, as others have done before (*22*). In yeast, expression of RHM1 and RHM1^G16A^, but not RHM1^A161K^, drove condensate formation (Fig. 1G). We measured accumulation of UDP-rhamnose by expressing RHM1 in yeast followed by extracting nucleotide-sugars and detecting UDP-rhamnose by liquid chromatography-mass spectrometry (LC-MS). The parent yeast strain W303a had no detectable UDP-rhamnose while W303a transiently expressing RHM1 made a substantial amount of UDP-rhamnose (Fig. 2J and Fig. S5). Unlike expressing RHM1 in W303a cells, UDP-rhamnose was undetectable in W303a cells expressing RHM1^A161K^ or RHM1^G16A^ (Fig. 2J). In line with our *in vivo* observations, this experiment suggests that RHM1 condensation is linked to its enzymatic activity.

### RHM1-bodies regulate additional UDP-rhamnose synthesis enzymes

We next wondered whether RHM1-bodies implicate other enzymes in the rhamnose synthesis pathway. Despite RHM1 being sufficient to synthesize rhamnose, the pathway also includes UDP-4-KETO-6-DEOXY-D-GLUCOSE-3,5-EPIMERASE-4-REDUCTASE 1 (UER1), which resembles a truncated version of RHM1 with only the BD2 domain being present. Therefore, UER1 cannot perform the initial catalytic step using UDP-glucose and can only synthesize UDP-rhamnose when the intermediate UDP-4-keto-6-deoxyglucose (UDP-4K6DG) is present (*4*) (*23*). To test if UER1 contributes to UDP-rhamnose synthesis *in planta*, we measured cell wall-associated rhamnose via neutral monosaccharide analysis in *uer1-1* seedlings. Plants carrying the *uer1-1* allele had reduced cell wall-associated rhamnose compared to Col-0 plants (Fig. 3A), confirming UER1 functions in the rhamnose synthesis pathway *in planta*. In addition, *rhm1-2;uer1-1* plants did not have a significant decrease in cell wall-associated rhamnose in seedlings compared to *rhm1-2* and *uer1-1* single mutants, consistent with UER1 requiring RHM1 for function *in planta* (Fig. 3A). Despite *rhm1-2;uer1-1* seedlings not having a significant decrease in cell wall rhamnose compared to single mutants, we observed *rhm1-2;uer1-1* had a more severe petal phenotype compared to either single mutant. Petals of *uer1-1* resemble the twisted *rhm1-2* petal phenotype, but are larger than *rhm1-2* (Fig. 3B). In the double *rhm1-2;uer1-1* mutant, flower petals are smaller than the *rhm1-2* and *uer1-1* single mutants (Fig. 3B), indicative of an additive genetic interaction between RHM1 and UER1 in petal tissues. Since UER1 lacks BD1, we speculated that UER1 could only function in UDP-rhamnose synthesis if UER1 is physically near RHM1-bodies. UER1 is predicted to interact with RHM1 *in silico* (Fig. S6). To test whether RHM1-bodies sequester UER1 and are required for UER1’s function, we probed for their physical interaction by coexpressing UER1 with RHM1 in yeast cells. UER1 was strongly recruited to RHM1-bodies (Fig. 3C). We next tested if UER1 is sufficient to form biomolecular condensates by expressing UER1 by itself or by expressing UER1 with RHM1^A161K^. In both cases, UER1 remained diffuse (Fig. 3C), demonstrating UER1 is a client but not a scaffold of RHM1 condensates. We next tested if UER1 enhances UDP-rhamnose synthesis when coexpressed with RHM1 in yeast. UER1 alone did not synthesize UDP-rhamnose (Fig. 3D). Coexpressing UER1 with RHM1 accumulated more UDP-rhamnose compared to expressing RHM1 alone (∼1.6 fold more; Fig. 3D), further suggesting an additive interaction between UER1 and RHM1. Together, we conclude that RHM1-bodies are required for UER1 condensation and the close proximity of UER1 to RHM1 through sequestration to RHM1-bodies is required for UER1 function, highlighting RHM1-bodies as an organizational unit that regulates the UDP-rhamnose synthesis pathway.

**Figure 3.**
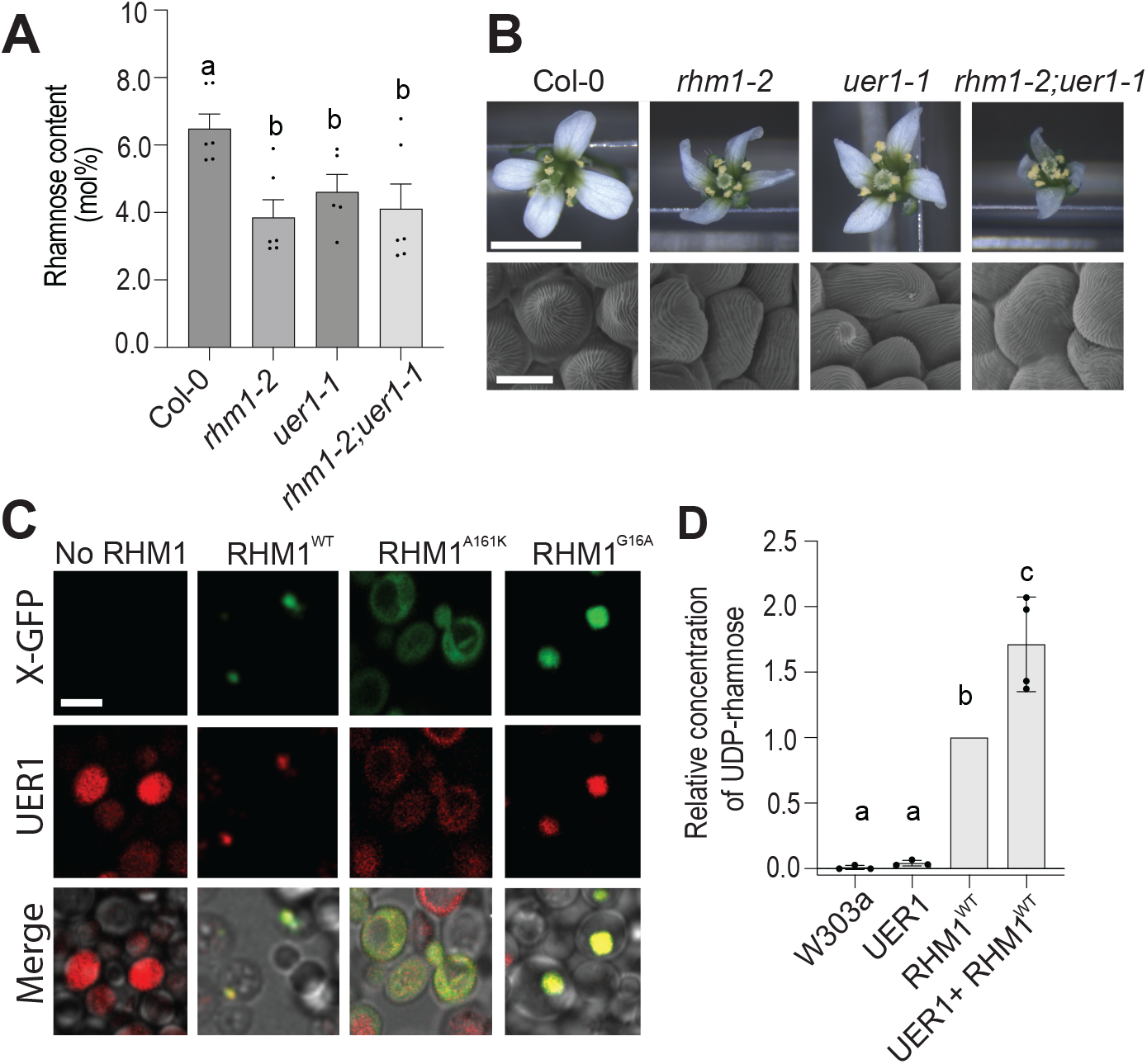
Rhamnosomes are required for UER1 to function in the UDP-rhamnose synthesis pathway. **A**) Quantification of rhamnose in the cell wall in seedlings of Col-0, *rhm1-2, uer1-1*, and *rhm1-2*; *uer1-1*. Data is from 6 biological replicates. For statistical significance, different letters represent significant differences (p-value < 0.05, ANOVA). **B**) Comparison of flowers from Col-0, *rhm1-2, uer1-1*, and *rhm1-2*; *uer1-1. Top*, flowers of Col-0, *rhm1-2, uer1-1*, and the *rhm1-2*; *uer1-1* plants (scale bar = 5 mm). *Bottom*, petal cone cells of the mutants show parallel rather than radial patterns of secondary cell wall deposition (scale bar = 10 μm). Micrographs are representative of multiple independent experiments. **C**) Subcellular localization of UER1 when coexpressed with RHM1 and RHM1 point mutants in yeast. *Top*, X-GFP fluorescence channel corresponds to No RHM1, RHM1-GFP, RHM1^A161K^-GFP, and RHM1G16A-GFP. *Middle*, UER1-RFP fluorescence. *Bottom*, merge of top and middle images with their corresponding brightfield micrograph. Micrographs are representative of 3 independent experiments. All micrographs are the same magnification (scale bar = 5 μm). **D**) Relative concentration of UDP-rhamnose in yeast cells transiently expressing UER1, RHM1, or RHM1 and UER1. Concentrations for all strains were normalized to the UDP-rhamnose concentration synthesized by RHM1 for each replicate. Data is from 3-4 samples, from 2 independent experiments. For statistical significance, different letters represent a p-value < 0.05 (ANOVA).

## Discussion and Conclusion

Biomolecular condensates have emerged as a major organizing principle in the cell (*24, 25*), yet evidence of whether condensates are essential in metabolic processes has been sparse. In the plant kingdom, the pyrenoid is an algal condensate that aids the inefficient enzymatic fixation of CO_2_ by concentrating substrate and enzyme (*26*), but additional examples of condensates involved in metabolism remain scarce. Further, the identification and function of biomolecular condensates formed by metabolic enzymes, and their impact on development in multicellular organisms, is a new frontier in biology (*25*). Here, we identified and characterized a novel biomolecular condensate in plants that functions in the synthesis of UDP-rhamnose, which is important in cell wall architecture and plant development. We name this RHM1-body the rhamnosome.

After identifying the rhamnosome as a novel condensate, and not a part of stress granules as previously reported (*12*), we probed its role in UDP-rhamnose synthesis. Throughout this study, we sought to identify if rhamnosomes are a site for storing inactive RHM1 enzymes or if they are associated with enhancing rhamnose synthesis. In this study, several lines of observation indicate that the rhamnosome forms to enhance UDP-rhamnose synthesis. First, we demonstrated that rhamnosomes form during developmental stages linked to increased cell wall synthesis. Second, rhamnosome formation was required for UDP-rhamnose synthesis in a heterologous system that does not make rhamnose. Third, UER1 localized to rhamnosomes and required rhamnosome formation for function in UDP-rhamnose synthesis. These multiple lines of evidence suggest that rhamnosome formation via RHM1 condensation is required to enhance, not inhibit, UDP-rhamnose synthesis. Together, our results constitute a novel intersection of condensates serving as important metabolic hubs for cellular metabolism, which are required for organismal development.

Rhamnosome formation being dynamically regulated during plant development poses a question of what controls their formation and dissolution. We observed that the frequency of cells with rhamnosomes in a tissue increases in stages which require UDP-rhamnose, followed by a decreased frequency in stages that do not require UDP-rhamnose. Early stages of petal development (stage 11) had high RHM1-GFP signal, but rhamnosomes did not form until stage 12. In contrast to rhamnosome formation between petal stages 11 and 12, RHM1 in heterologous systems constitutively formed rhamnosomes, suggesting there are plant factors that can prevent RHM1 condensation. For example, the SG proteins CAPRIN and USP10 interact with the SG scaffold protein G3BP to regulate formation and dissolution of SGs (*27*). Similar proteins may exist in plants to regulate rhamnosome condensation. In contrast to the increased frequency of rhamnosome formation during petal developmental stages 11 and 12, between flower stage 13 and stage 14 we observed the frequency of rhamnosomes in petal cells decreased. The mechanism of rhamnosome dissolution remains an open question. We currently do not know if rhamnosomes are degraded or disassembled. Investigating other signaling cues that intersect with rhamnosome formation and dissolution will identify additional regulatory circuits responsible for rhamnosome regulation during plant development. If future work can identify novel plant-specific mechanisms of regulating RHM1 condensation and dissolution, the principles governing formation and function can be engineered into heterologous systems to generate both novel condensates and engineer molecular circuits to regulate their formation.

The discovery of rhamnosomes opens new lines of inquiry. How deeply are rhamnosomes conserved and when did they evolve? Outside of plants, rhamnose is synthesized in bacteria (*28, 29*), fungi (*30*), protozoa (*31*), and some invertebrate animals (*32*). RHM1 homologs (composed of both BD1 and BD2) are present in unicellular algae, mosses, and angiosperms (Fig. S7A). Heterologous expression of RHM1 from the unicellular algae *Ostreococcus tauri* formed rhamnosomes (Fig. S7A and B), suggesting rhamnosomes are formed throughout the plant kingdom. In prokaryotes and non-plant eukaryotes that synthesize rhamnose, such as *Escherichia coli, Trichomonas vaginalis*, and *Caenorhabditis elegans*, nucleotide diphosphate (NDP)-rhamnose synthesis typically requires three monofunctional enzymes (*23, 31, 32*). *E. coli* requires rhamnose for synthesis of glycan used for the lipopolysaccharide in the cell wall (*33*) and the first monofunctional enzyme in rhamnose synthesis is a member of the NAD(P)-binding domain superfamily (dTDP-glucose 4,6-dehydratase 2; rffG) (*34*). rffG has conserved residues with BD1 in the helix associated with RHM1 condensation (region around A161; Fig. S7C), suggesting rffG may form condensates. Similarly to *E*.*coli*, the protozoan *T. vaginalis* synthesizes rhamnose for their lipoglycan outer surface which is an important virulence factor (*31*). *Trichomonas vaginalis* NDP-D-Glucose 4,6-dehydratase forms homodimers (*31*) and is similar in protein sequence to RHM1 BD1. In the multicellular worm *C. elegans*, rhamnose is required for embryogenesis and larval molting, likely by forming a component of the outer cuticle (*32*). *C. elegans* rhamnose synthesis also requires the monofunctional protein NDP-D-Glucose 4,6-dehydratase (RML-2) (*32*), which also shares conserved residues with the BD1 helix associated with RHM1 condensation in *Arabidopsis* (Fig. S7C). Together, rhamnose in prokaryotes and eukaryotes appears to be an important component of the outer cell layer. Based on the current literature and our study, compartmentalization of the rhamnose pathway may serve as a universal cellular method to regulate rhamnose biosynthesis throughout life.

Rhamnosomes shine a new light on how cells use biomolecular condensates to regulate their metabolism during organismal development. Rhamnosomes can be constituted from a single protein and are functional in heterologous systems, making these organelles a novel platform to study how condensation impacts enzymatic function. Together, the molecular rules and design principles gleaned from this and future studies on the rhamnosome may inspire the biocatalysts of tomorrow that are increasingly needed in building a green economy.

## Acknowledgments

We would like to thank members of the Rhee lab for helpful discussion on this project. We would like to thank Andrey Malkovskiy at the Carnegie Institution for Science Microscopy Core for assistance with microscopy. We would also like to thank Dan Jones, Anthony Schilmiller, and Lijun Chen at the Research Technology Support Facility (RTSF) Mass Spectrometry and Metabolomics Core at Michigan State University for help in the analysis of UDP-rhamnose. We would also like to thank the Vivian Irish lab (Yale University) for providing the *Arabidopsis rhm1-2* line. This work was done in part on the ancestral land of the Muwekma Ohlone Tribe which was and continues to be of great importance to the Ohlone people, and on the ancestral, traditional, and contemporary Lands of the Anishinaabeg – Three Fires Confederacy of Ojibwe, Odawa, and Potawatomi peoples. Dedicated in the memory of Dennis Skupinski.

## Author contributions

Conceptualization: SF, SYR, YD, SB, WPD

Methodology: SF, SYR,YD, SB, WPD

Investigation: SF, WPD, JAC, RH, SB, YD, IW, MB, OMSC, DR, JF

Visualization: SF, JAC, RH, SB, YD, OMSC, DR, JF

Funding acquisition: SYR, SB, DR, JF, MB

Project administration: SYR

Supervision: SF, SYR

Writing – original draft: SF, WPD, SYR

Writing – review & editing: SF, SYR, YD, SB

## Data and materials availability

All data are available in the main text or the supplementary materials.

## Supplementary Materials

### Materials and Methods

#### Molecular Cloning

The primers and plasmids used in this study are available in Tables S1 and S2. Codon optimized constructs are listed in Table S3.

### Constructs for transient expression in Nicotiana benthamiana

Constitutively expressed *p35S*::RHM1-GFP and *p35S*::UBP1c-RFP constructs were generated by first amplifying the RHM1 and UBP1c coding sequence from Col-0 cDNA using PCR. RHM1 and UBP1c amplicons were recombined by Gateway cloning (Gateway BP Clonase II, ThermoFisher) into pDONR221. Both RHM1/pDONR221 and UBP1c/pDONR221 were cloned into pGWB605 and pGWB660 (*35*), respectively, using Gateway LR Clonase II (ThermoFisher).

### Constructs for stable expression in Arabidopsis lines

Constructs for stable expression in *Arabidopsis* lines were generated through custom synthesis and subcloning by Genscript. The native *Arabidopsis* RHM1 promoter comprising the genomic DNA 1.5 kb upstream of the RHM1 translation start site, along with the *Arabidopsis* RHM1 coding sequence, were synthesized and subcloned into the pGWB604 vector (*35*) by GenScript (Piscataway, USA). Both *pRHM1:*:RHM1^A161K^/pGWB604 and *pRHM1:*:RHM1^G16A^/pGWB604 were generated from *pRHM1:*:RHM1/pGWB604 using site-directed mutagenesis (GenScript).

### Constructs for expression in Saccharomyces cerevisiae

Constructs for yeast expression were generated through custom synthesis and subcloned into pESC-URA and pESC-Trp vectors by Genscript. To generate constructs for expressing RHM1 in yeast, the DNA sequence for *Arabidopsis* RHM1 was synthesized in-frame with an N-terminal GFP fusion by GenScript. This RHM1-GFP construct was then subcloned into the pESC-URA plasmid (GenScript). Both RHM1^A161K^/pESC-URA and RHM1^G16A^ /pESC-URA were synthesized by site-directed mutagenesis from the RHM1-GFP/pESC-URA (GenScript). A similar approach was used to generate a yeast codon optimized UER1 with a C-terminal mCherry tag, which was synthesized and subcloned into pESC-Trp (GenScript).

### Constructs for expression in human cell lines

Constructs for expressing RHM1 and G3BP in human cell lines were generated through custom synthesis and subcloned into a pcDNA3.1 backbone (GenScript).

## Plant system techniques and assays

### Plant Growth conditions

*Arabidopsis thaliana* and *Nicotiana benthamiana* plants were grown under long day conditions (16 hours (hrs) light: 8 hrs dark, temperature of 21°C, humidity ∼50 %, and a light intensity of 150 μmol/m^2^).

### Plant Transformation

A list of plant lines used in this study are available in Table S4. *Arabidopsis* lines were generated by transforming plants carrying the *rhm1-2* allele (a kind gift from the Irish Lab, Yale University) with the *pRHM1:*:RHM1/pGWB604, *pRHM1:*:RHM1^A161K^/pGWB604, or *pRHM1:*:RHM1^G16A^/pGWB604 construct. Transient transfection of *Nicotiana benthamiana* was performed as described in (*36*).

*Arabidopsis transformation. rhm1-2* plants were transformed using the floral dip method (*37*). Briefly, *Agrobacterium tumefaciens* GV3101 carrying the desired construct, described above, were grown overnight, and cells were pelleted at 4,000 x g for 10 minutes. Pelleted cells were resuspended to an OD_600_ of 1.0 in resuspension solution (5 % (w/v) sucrose and 0.05 % (v/v) Silwet L-77 (VIS-01, Vac-In-Stuff)). Flowering *Arabidopsis* plants were inverted and the above-ground tissue was dipped into the resuspended *A. tumefaciens* solution for 10 sec. Plants were incubated in a dark environment at room temperature overnight, and the following day, the plants were transferred to a growth chamber and grown in long day conditions, as described above. *Arabidopsis* seeds transformed with the transgenes were selected on ½ x MS media with 25 μg/mL glufosinate. Herbicide-resistant seedlings were transferred to soil and were grown under long day conditions.

### Plant stress conditions for *Nicotiana* leaves

To test if heat, UV, or oxidative stress cause RHM1 to condense in transiently transfected *Nicotiana* leaves, whole *Nicotiana* leaves were heat stressed by exposing them to 42°C for 1 hour (hr). Oxidative stress was induced by incubating whole leaves in 1 % H_2_O_2_ for 1 hr. UV stress was induced by treating whole leaves with 1500 μJ of 254 nm UV light (UV Stratalinker 1800, Stratagene) and imaged 1 hr later.

### Plant phenotype assays

All plants for an independent experiment were grown together on soil in the same flat, which was composed of 4 - 6 plants for each genotype.

*Flower petal imaging and measurement*. Flower petals were imaged using a Leica M165 FluoCombi (FC) microscope. For comparing flower phenotypes between genotypes, all plants were grown together, and images were taken on the same day from stage 14 flowers from all genotypes. For measuring petal area, images of flowers were analyzed using the built-in area measurement tool in ImageJ version 1.54.

*Measuring seed mucilage*. Seed mucilage depth was measured by a method similar to (*21*). First, seeds were hydrated for 1 hr in water with 0.1 % (w/v) ruthenium red (Sigma, CAS-No 11103-72-3). After 1 hr, the outer layer of mucilage was removed using mechanical stress generated by vortexing the seeds for 60 sec, in order to allow analysis of the adherent mucilage layer. The seeds were then washed 3 times with water, and the adherent mucilage layer was imaged using a Leica M165 FC microscope. Images of seeds stained with ruthenium red were analyzed using the built-in measurement tools in ImageJ.

*Rosette area measurements*. Plants for rosette area measurements were thinned out to one plant per pot after ∼1 week. Whole flats were imaged at week 3, and these images were used to calculate rosette area using the built-in area measurement tool in ImageJ.

*Neutral monosaccharide cell wall analysis. Arabidopsis* seeds were planted and grown under long days (16 hr light/ 8 hr dark) at 22°C for 7 days on MS-agar (½ x MS salts, 10 mM MES-KOH pH 5.7, 1 % (w/v) phytoagar, 1 % (w/v) sucrose). Seedlings were harvested and incubated in 10 mL of 70 % (v/v) ethanol for 6 hr at room temperature with gentle shaking. The supernatant was removed and replaced with 10 mL of 70 % (v/v) ethanol followed by incubation for 18 hr at 25°C. The supernatant was removed, and seedlings were incubated with 10 mL of 1:1 chloroform: methanol for 24 hr at 25°C. The supernatant was removed, seedlings were transferred to pre-weighed 2 mL screw cap tubes, and dried for 24 hr. Dried seedling tissue was milled with steel balls in a Retsch mill MM300 at 25 Hz for 2.5 minutes to produce ball-milled alcohol insoluble residue (AIR). Neutral monosaccharides were released from 3-5 mg of AIR, derivatized to alditol acetates, and analyzed by gas chromatography as previously described (*38*).

### Microscopy for plant systems

*Imaging subcellular bodies*. RHM1-GFP was visualized using a Leica SP8 scanning laser confocal microscope (Leica, Wetzlar, Germany). Proteins tagged with GFP were excited with a white light laser set to 488 nm, and fluorescence was detected using a hybrid detector for light in the range of 498 nm - 546 nm. Proteins tagged with mCherry were excited with a white light laser at 561 nm and fluorescence was detected in the range of 574 nm - 629 nm by a hybrid detector. For *Arabidopsis* tissues, all cells within an area 100 μm x 100 μm were imaged. For rhamnosome quantification and 3D projections, all cells within 100 μm x 100 μm x 30 μm were imaged. A similar approach was used for imaging yeast, with the exception that cells were only imaged in the x and y dimension.

*Visual presentation of micrographs*. Figure images were processed using LasX software (version 3.7.4), or Imaris x64 (version 9.8.0) for 3D images. Micrographs were directly exported from LasX and used to generate figures. For 3D reconstruction of SG and rhamnosomes, 3D reconstruction was performed on a set of images composed of the x, y, and z planes, generated as described above. Files were exported from the Leica microscope software LasX (exported as a .lif file) and converted into .ims files using Imaris File Converter. Tobacco cell images (in the x, y, and z planes) were used to generate 3D reconstructions of GFP-marked rhamnosomes and UBP1c-RFP-marked stress granules in Imaris x64 using the built-in ‘Surface’ algorithm set to 0.25 μm surface detail.

*Quantification of rhamnosomes*. Quantifying the number of rhamnosomes per tissue section was performed similar to quantification of other cytoplasmic granules described in (*36*). Briefly, the number of cells with rhamnosome in a tissue was measured by generating sets of images in x, y, and z dimensions. Micrographs were then divided into subsections composed of 40 μm x 40 μm x 20 μm and projected into a single plane. For quantifying the number of cells with rhamnosomes, the total number of cells in the image was compared to the number of cells with rhamnosomes.

*Imaging nanoridges of petal cone cells*. Images of nanoridges on petal cone cells were obtained using a Scanning Electron Microscope (Quanta 200 SEM, Thermofisher, Waltham, MA, USA) equipped with Back Scatter Electron (BSE) detectors in high-vacuum mode. Tissue was prepared by fixing whole flowers in 100 % methanol for 2 hr. Methanol was decanted, replaced with 100 % ethanol and incubated for 3 hr, then decanted and replaced with fresh 100 % ethanol. The tissue was desiccated using a DCP1 Critical Point Drying Apparatus (Denton Vacuum) with liquid carbon dioxide. Once samples were desiccated, the dried samples were sputtered with gold (15 % for 3 minutes) using a Desk IV Sputter Coater (Denton Vacuum), and imaged on the SEM.

### Immunoblotting for RHM1-GFP expression

RHM1-GFP protein was detected by Western blotting with a protocol modified from (*36*). Briefly, all above ground tissue of 3-week-old plants was collected and frozen in liquid nitrogen. Tissue was pulverized and the resulting powder was resuspended in 500 μL of chilled extraction buffer (50 mM Tris–HCl pH 7.5, 0.15 M NaCl, 10 % (v/v) glycerol, 0.01 % (v/v) NP-40, 1 mM dithiothreitol, protease inhibitor cocktail (Roche, EDTA free, one tablet per 10 mL of resuspension buffer). Samples were incubated on ice for 5 minutes, and then centrifuged for 10 minutes at 16,000 x g at 4°C. The total amount of protein was measured using a Bradford assay (Pierce Bradford Plus Protein Assay; Thermo Scientific), and 30 μg of total protein from the soluble fraction was resolved on a 12 % w/v SDS-PAGE gel (Mini-Protean TGX, Bio-Rad). The gel was then electroblotted onto a polyvinylidene fluoride (PVDF) membrane using a wet transfer method. The membrane was blocked with 3 % dry milk in 1 x phosphate buffer saline with 0.1 % Tween 20 (PBST) for 1 hr, and immediately incubated with polyclonal rabbit anti-GFP IgG (AS20 4512; Agrisera) for 1 hr at room temperature. The membrane was washed 5 x in 1 x PBST, followed by incubation with horseradish peroxidase coupled goat anti-rabbit IgG (AS09 602; Agrisera). The membrane was washed 3 x in PBST followed by 2 x in PBS. Chemiluminescent detection was performed using the Super Signal West Dura Extended Duration Substrate kit (ThermoFisher).

## Human cell culture system assays

### Human cell culture growth and microscopy

U2OS cells (ATCC, HTB-96) were grown at 37°C in a humidified atmosphere with 5 % CO_2_ for 24 hr in Dulbecco’s Modified Eagle’s Medium (DMEM), high glucose (4500 mg/L glucose), and 1 x GlutaMAX + 10 % Fetal Bovine Serum (FBS). Cells were transiently transfected using Lipofectamine 3000 (Thermo Fisher Scientific) according to manufacturer’s instructions. 24 hr after transfection, cells grown on coverslips were fixed in 4 % formaldehyde in PBS for 20 minutes at room temperature. Nuclei were stained with Hoechst 33258 (Life Technologies) at 1 μg/mL in PBS for 10 minutes at room temperature. Coverslips were then washed with PBS and mounted on slides using ProLong Gold antifade (Life Technologies). Confocal images were obtained using a Zeiss laser scanning microscope (LSM) 900 confocal 40x objective.

### Human cell culture stress treatments

To induce stress granule formation, cells were treated for 1 hr with 500 μM of sodium arsenite in PBS (Sigma-Aldrich). To visualize stress granules, fixed coverslips were incubated in blocking buffer (5 % goat serum, 0.5 % BSA, 0.4 % TritonX100 in PBS) for 1 hr at room temperature, then incubated with anti-G3BP1 antibody (Abcam ab181150, 0.306 mg/mL diluted 1:500) overnight at 4°C. After 3 x washes with 0.4 % TritonX100 in PBS, coverslips were incubated with goat anti-rabbit IgG conjugated to AlexaFluor-594 (Life Technologies, diluted 1:500) for 80 minutes at room temperature. Coverslips were stained with Hoechst and mounted as mentioned above. All stains and antibodies for immunofluorescence assays were diluted in the blocking buffer.

### Image analysis of human cell lines

Representative cell subsections and line-scan measurements were collected using FIJI (version 1.54). Line-scan gray values were smoothed with a Savitzky-Golay filter (scipy 1.11.3) and plotted using matplotlib (3.7.2).

## Yeast system assays

### Yeast cell growth and transformation

*Saccharomyces cerevisiae* (strain W303a; ATCC 208352) cells were grown in Difco YPD Broth (#242820). To transform W303a cells with expression constructs, W303a cells were transformed with RHM1/pESC-URA, RHM1^A161K^/pESC-URA, RHM1^G16A^/pESC-URA, or UER1/pESC-TRP following the Frozen EZ Yeast Transformation II Kit protocol (T2001, Zymo). Cells were grown in synthetically defined (SD) media without URA (for pESC-URA constructs) or without TRP (for pESC-TRP). To coexpress UER1, yeast cells with RHM1/pESC-URA, RHM1^A161K^/pESC-URA, or RHM1^G16A^/pESC-URA were transformed using the Frozen EZ Yeast Transformation II Kit with UER1/pESC-Trp, and grown on SD media without URA/TRP. SD media was supplemented with 1 x yeast nitrogen base without amino acids and ammonium sulfate (Difco #233520) and 2 % dextrose, or by replacing dextrose with 2 % galactose (final concentration; Sigma, CAS-No 59-23-4) to induce protein expression.

### Microscopy on yeast cells

Microscopy on fluorescently tagged proteins in yeast used the same parameters and techniques as for microscopy in plants (described above). For quantifying the frequency of RHM1-body formation in yeast, RHM1/pESC-URA, RHM1^A161K^/pESC-URA, RHM1^G16A^/pESC-URA, and UER1/pESC-TRP were induced with 2 % galactose for 24 hr prior to imaging.

### Growing yeast for UDP-rhamnose assays

For UDP-rhamnose synthesis assays, yeast cells were grown overnight at 28°C in 10 mL of SD media with 2 % dextrose and 1 x yeast nitrogen base (as described above). After 24 hr, 1 mL of the overnight culture was used to inoculate 200 mL SD media with 2 % galactose and 1 x nitrogen base. The 200 mL cultures were grown at 28°C for 48 hr. The cell density of each culture was measured prior to centrifugation and used to normalize downstream LC-MS/MS data (further described below). Cells were collected by centrifugation of cultures at 4000 x g, followed by discarding the supernatant, and freezing the resulting cell pellet until nucleotide sugar extraction, which is described below.

### Measuring UDP-rhamnose in yeast cells

For nucleotide sugar extraction and analysis using yeast cells, we used a modified version of the method presented in (*22*). Briefly, pelleted yeast cells were resuspended in 5 mL of 75 % methanol containing 0.1 % formic acid, centrifuged, and the supernatant was collected and evaporated until dry. The samples were reconstituted in 1 mL 80 % acetonitrile. UDP-rhamnose was measured by LC-MS at the Research Technology Support Facility (RTSF) Mass Spectrometry and Metabolomics Core at Michigan State University. Samples used cyclic dimeric guanosine monophosphate (C-di-GMPF) as an internal standard. UDP-rhamnose concentrations were determined by measuring the integral of the LC-MS peak relative to the internal standard.

### *In silico* modeling

#### Structural analysis of RHM1

The predicted structure of RHM1 (At1g78570) was generated using the AlphaFold protein structure database (UniProt ID: Q9SYM5) (*39*). Protein structure figures were created using either Chimera (version 1.16) (*40*) or PyMOL (version 2.5).

#### Computational modeling of the RHM1 protein and dimerization

Molecular modeling, alignments, and analyses were performed with UCSF Chimera (version 1.16) (*39*) and ChimeraX (version 1.6.1) (*41*). RHM1 mutant structural prediction and RHM1-UER1 protein-protein interaction were generated using ChimeraX (version 1.6.1)(*39*) via ColabFold cloud computing (Python 3, V100 GPU), an AlphaFold-based Google notebook (*42*).

#### Phylogenetic tree construction

Protein sequences for RHM1-homologs from *viridiplantae* proteins were identified using NCBI blast. Protein sequences from bacteria (*28, 29*), fungi (*30*), protozoa (*31*), and invertebrate animals (*32*) were identified from previous literature, and their amino acid sequences were retrieved from Uniprot. Protein sequences used for alignment and phylogenetic tree construction are in Table S5. Protein alignment used Geneious (version 2023.2) and the built in Clustal Omega sequence alignment algorithm using default settings. Phylogenetic tree construction and consensus sequence visualization used default settings in Geneious.

#### Statistics

Unless otherwise noted, t-tests were used for statistics comparing two samples, and ANOVA was used to compare multiple samples. Graphs and statistical tests used Prism (GraphPad; version 9.3.1).

#### Data presentation

Unless otherwise noted, all graphs present the standard deviation of the data. For statistical significance, different letters represent a p-value < 0.05.

**Supplementary Figures. S1 to S7**

**Supplementary Figure 1**.

Rhamnosomes are distinct from stress granules.

**Supplementary Figure 2**.

RHM1 dimerization requires hydrophobic interactions between RHM1 monomers.

**Supplementary Figure 3**.

Characterization and comparison of RHM1-GFP complementation lines.

**Supplementary Figure 4**.

Complementation of novel *rhm1-2* phenotypes.

**Supplementary Figure 5**.

RHM1 is sufficient to synthesize UDP-rhamnose in yeast.

**Supplementary Figure 6**.

*In silico* prediction of RHM1-UER1 interaction.

**Supplementary Figure 7**.

Evolution of RHM1 and rhamnosomes.

**Tables S1 to S4**.

**Table S1**.

Primers used in this study.

**Table S2**.

Plasmids used and generated in this study.

**Table S3**.

Sequences for codon optimized expression constructs.

**Table S4**.

Plant lines used and generated in this study.

**Table S5**.

Protein sequences of RHM1 homolog proteins used for phylogenetic analysis.

