## Supplemental Tables for "*Arabidopsis thaliana* RHAMNOSE 1 condensate formation drives UDP-rhamnose synthesis"

**The PDF file includes:**

Tables S1 to S5

### Supplementary Tables

**Table S1.** Primers used in this study.

| **Primer name** | **5'- 3' primer sequence** | **notes** |
|---|---|---|
| pricdsG3BP-FWD | GGGGACAAGTTTGTA CAAAAAAGCAGGCTTCATGGCGACTCCTTAT CCTGG | CDS cloning of *G3BP1* without the stop codon |
| pricdsG3BP-REV | GGGGACCACTTTGTA CAAGAAAGCTGGGTC GCGACCACCACCGCGG TAGT | CDS cloning of *G3BP1* without the stop codon |
| pricdsRHM1-FWD | GGGGACAAGTTTGTA CAAAAAAGCAGGCTT CATGGCTTCGTACACT CCCAA | CDS cloning of *RHM1* without the stop codon |
| pricdsRHM1-REV | GGGGACCACTTTGTA CAAGAAAGCTGGGTC GGTTTTCTTGTTTGGCC CGT | CDS cloning of *RHM1* without the stop codon |
| SF044 F | ACACTCCCAAGAACATTCTC | Genotyping *rhm1-2* |
| SF045 R | TTGTTTGGCCCGTATGCATA | Genotyping *rhm1-2* |
| WPD_UER1_F | TCATTTAACTTCCTAATCTA | Genotyping *uer1-1* |
| WPD_UER1_R | TTCCTCCAATGTGAAGTTCTT | Genotyping *uer1-1* |

**Table S2.** Plasmids used and generated in this study.

| Name in paper | Protein | Tag | Plasmid | Exp |
|---|---|---|---|---|
| RHM1-GFP | RHM1 | GFP | pGWB605 | Nb expression |
| UBP1c-RFP | UBP1c | RFP | pGWB660 | Nb expression |
| RHM1-GFP | RHM1 | GFP | pGWB604 | At expression |
| $RHM1^{A161K}$-GFP | $RHM1^{A161K}$ | GFP | pGWB604 | At expression |
| $RHM1^{G18A}$-GFP | $RHM1^{G18A}$ | GFP | pGWB604 | At expression |
| RHM1-GFP | RHM1 | GFP | pESC-URA | Yeast expression |
| $RHM1^{A161K}$-GFP | $RHM1^{A161K}$ | GFP | pESC-URA | Yeast expression |
| $RHM1^{G18A}$-GFP | $RHM1^{G18A}$ | GFP | pESC-URA | Yeast expression |
| UER1 | UER1 | mCherry | pESC-TRP | Yeast expression |

**Table S3.** Sequences for codon optimized expression constructs.

(Provided as .xslx file)

**Table S4.** Plant lines used and generated in this study.

| Name in paper | identifier/other names | Protein | Tag | Plasmid |
|---|---|---|---|---|
| Col-0 | Col-0 | n/a | n/a | n/a |
| *rhm1-2* | *rol1-2*, CS16373 | n/a | n/a | n/a |
| RHM1 | n/a | RHM1 | GFP | pGWB604 |
| RHM1$^{A161K}$ | n/a | RHM1$^{A161K}$ | GFP | pGWB604 |
| RHM1$^{G18A}$ | n/a | RHM1$^{G18A}$ | GFP | pGWB604 |
| *uer1-1* | CS926928 | n/a | n/a | n/a |

**Table S5.** Protein sequences of RHM1 homolog proteins used for phylogenetic analysis.

(Provided as .xslx file)
