## Supplemental Figures for "*Arabidopsis thaliana* RHAMNOSE 1 condensate formation drives UDP-rhamnose synthesis"

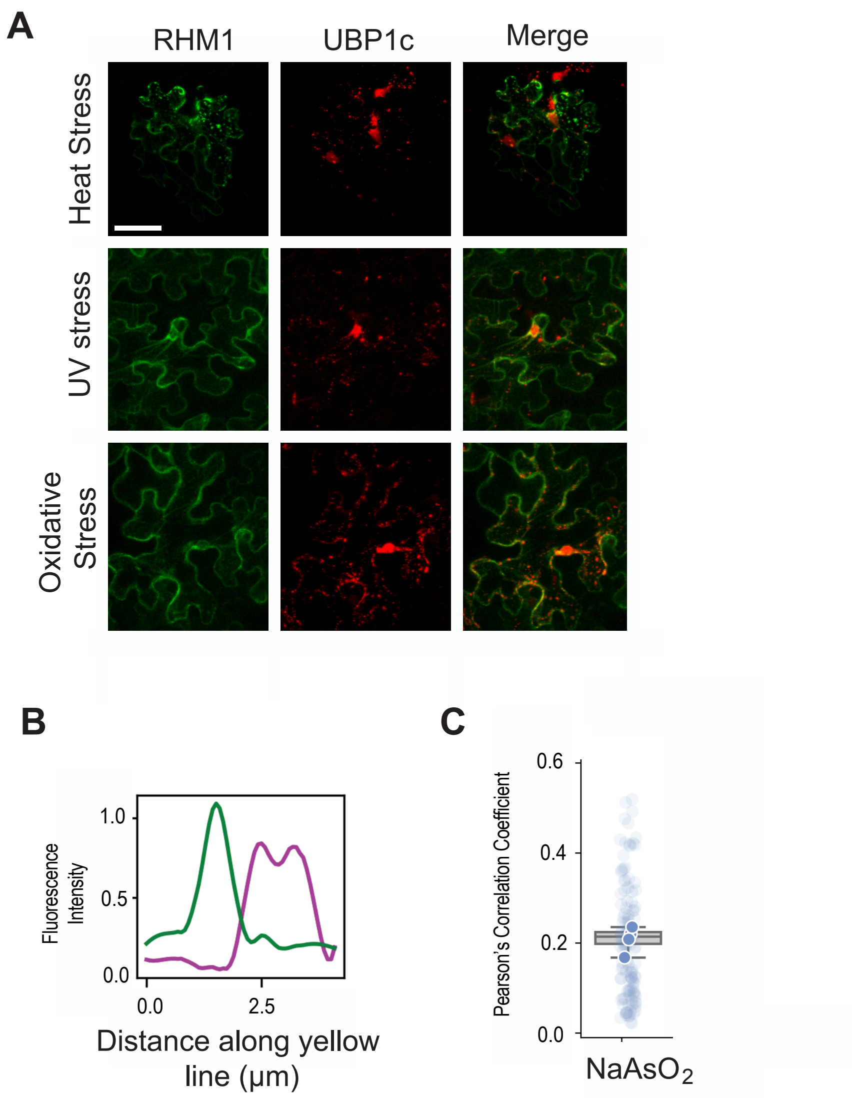


**Supplementary Figure 1. Rhamnosomes are distinct from stress granules.** **A**) Confocal images of transient transfection of *Nicotiana benthamiana* epidermal leaf cells expressing RHM1-GFP and the *Arabidopsis* stress granule marker UBP1c-RFP, challenged with various stresses (scale bar is 40 µm, all images are the same magnification). Stress conditions: heat stress induced by 1 hr at 42°C, oxidative stress induced by incubating leaf tissue in 1% $H_2O_2$ for 1 hr, UV stress by treating leaves with 1500 µJ of 254 nm UV light and imaging 1 hr later (see materials section for more details). Representative of 3 independent experiments. **B**) Fluorescent intensity of immunofluorescent anti-G3BP1 marked stress granule (magenta) and rhamnosome (green), X-axis corresponds to the length of the yellow line from Fig 1D. Curves were smoothed with a Savitzky-Golay filter. **C**) Quantification of colocalization between rhamnosomes and G3BP marked stress granules in human U2OS cells treated with sodium arsenite ($NaAsO_2$), using Pearson's correlation. A coefficient of 1 is perfect colocalization. Graph is from multiple cells from 4 independent experiments.

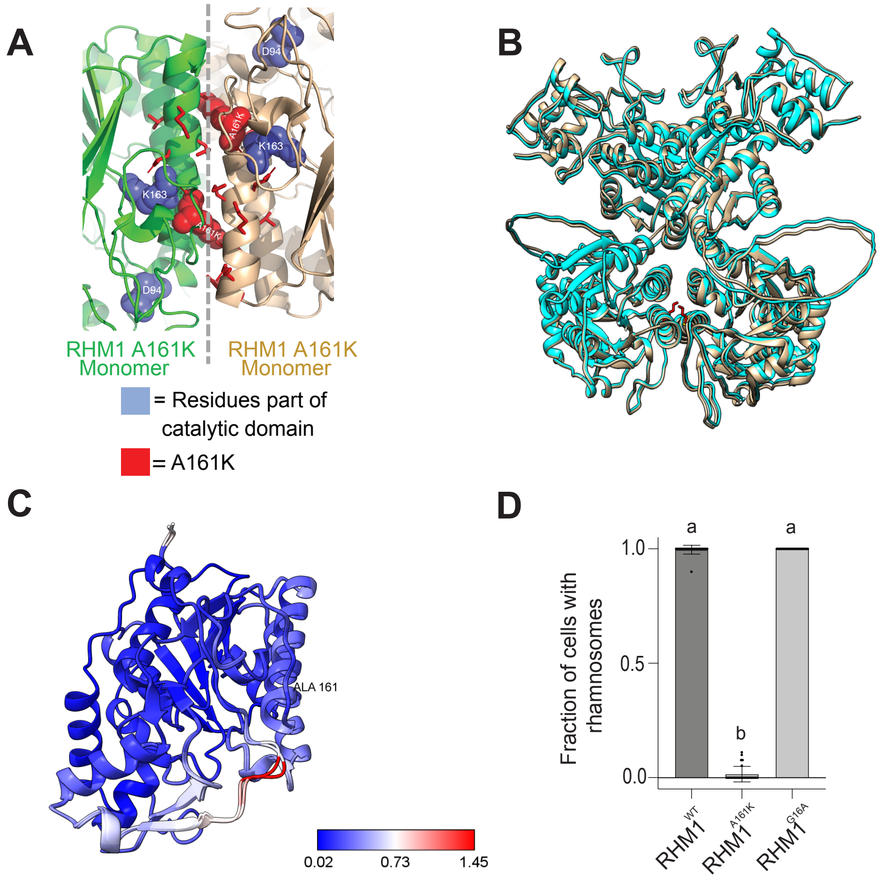


**Supplementary Figure 2. RHM1 dimerization requires hydrophobic interactions between RHM1 monomers. A**) Predicted dimerization for two RHM1$^{A161K}$ monomers. **B**) *In silico* prediction of RHM1 dimerization and protein structural alignment. RHM1 and RHM1$^{A161K}$ are superimposed, and the point mutant residue A161K is shown in red. Ribbon model is composed of RHM1$^{WT}$ (tan) and RHM1$^{A161K}$ (cyan). **C**) *In silico* prediction of RMSD of the BD1 for RHM1 and RHM1$^{A161K}$. Blue are RMSD values approaching 0.02 and red are values approaching 1.45. An RMSD value of 0 is perfect alignment of two proteins. **D**) Fraction of yeast cells with rhamnosomes. Yeast cells expressed either RHM1, RHM1$^{A161K}$, or RHM1$^{G16A}$. Data is from 3 independent experiments. For statistical significance, different letters represent a p-value < 0.05 (ANOVA).

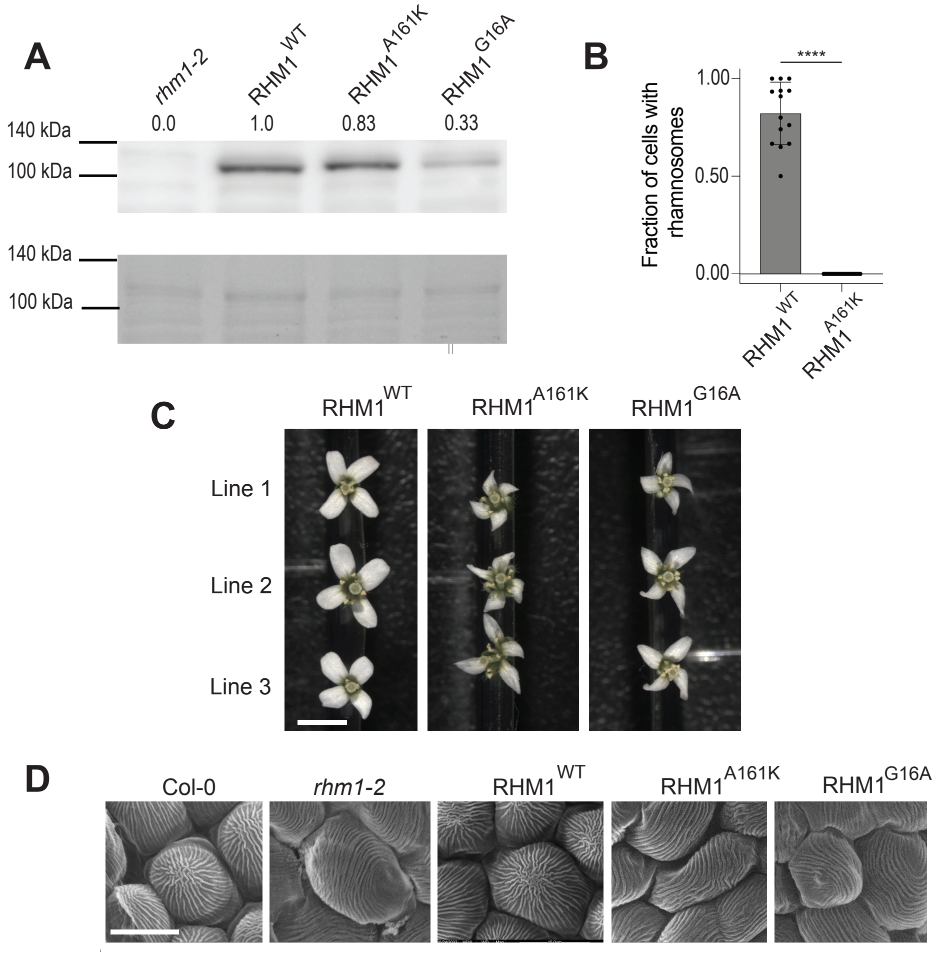


**Supplementary Figure 3. Characterization and comparison of the RHM1-GFP complementation lines. A**) Western blot comparing protein expression of *rhm1-2*, and the three *rhm1-2* lines complemented with *RHM1*::RHM1-GFP, *RHM1*::RHM1$^{A161K}$-GFP, and *RHM1*::RHM1$^{G16A}$-GFP. RHM1-GFP, RHM1$^{A161K}$-GFP, and RHM1$^{G16A}$-GFP are from the complementation lines from Figure 2. *Top*, western blot with the relative intensities of the protein listed above the image normalized to RHM1$^{WT}$-GFP. *Bottom*, PonceauS stain of the corresponding section of the membrane. Western blot used 30 µg of total protein, extracted from 3 week old *Arabidopsis* rosettes. RHM1-GFP is predicted to be 102.57 kDa. **B**) Fraction of cells with rhamnosomes in stage 13 petal cone cells. Each data point is the fraction of cells with rhamnosomes in a 40 µm x 40 µm section of petal tissue. Data is from 13-15 biological replicates from 3 independent experiments. For statistical significance, $p$-value < 0.05 (t-test). **C**) Comparison of *rhm1-2* flower complementation from three independent stable transgenic lines from each of the following constructs: *RHM1*::RHM1-GFP, *RHM1*::RHM1$^{A161K}$-GFP, and *RHM1*::RHM1$^{G16A}$-GFP. Plants were grown together and imaged together. Scale bar is 5 mm. **D**) Comparison of nanoridges on the surface of petal cone cells. Lines are the same as in Fig 2C. Scale bar is 10 µm.

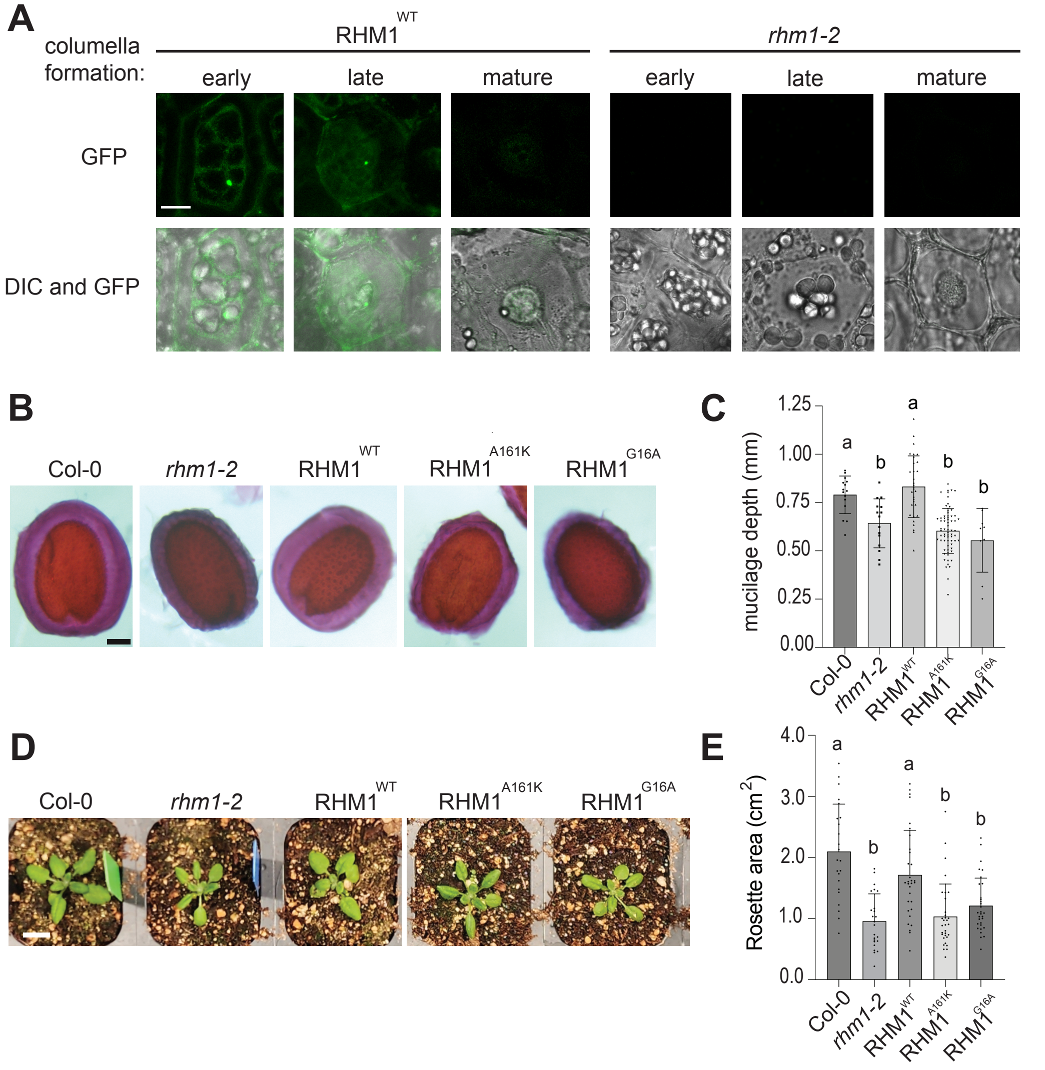


**Supplementary Figure 4. Complementation of novel *rhm1-2* phenotypes.**
**A**) Confocal micrographs of rhamnosome formation during stages of seed coat cell development. The same microscopy settings were used for RHM1-GFP in the *rhm1-2* background and *rhm1-2* (used as a control for seed autofluorescence). Scale bar is 10 µm, and all images are same magnification. **B**) Images of the adhesive inner mucilage layer of Col-0, *rhm1-2*, *RHM1*::RHM1-GFP, *RHM1*::RHM1$^{A161K}$-GFP, and *RHM1*::RHM1$^{G16A}$-GFP seeds. Parent plants were grown and harvested together. Mucilage is stained with ruthenium red. Scale bar is 0.1 mm. **C**) Quantification of the depth of the adhesive inner mucilage layer. Each data point is the mucilage depth of 1 seed. Data is from 15-30 seeds, from 3 independent experiments. For statistical significance, different letters represent a p-value < 0.05 (ANOVA). **D**) Image of rosette areas of 3 week old *Arabidopsis* Col-0, *rhm1-2*, *RHM1*::RHM1-GFP, *RHM1*::RHM1$^{A161K}$-GFP, and *RHM1*::RHM1$^{G16A}$-GFP plants. Plants were grown together and imaged together. Scale bar is 1 cm. **E**) Quantification of the total rosette area for the lines Col-0, *rhm1-2*, *RHM1*::RHM1-GFP, *RHM1*::RHM1$^{A161K}$-GFP, and *RHM1*::RHM1$^{G16A}$-GFP. Data is from 18-24 plants from 4 independent experiments. For statistical significance, different letters represent a p-value < 0.05 (ANOVA).

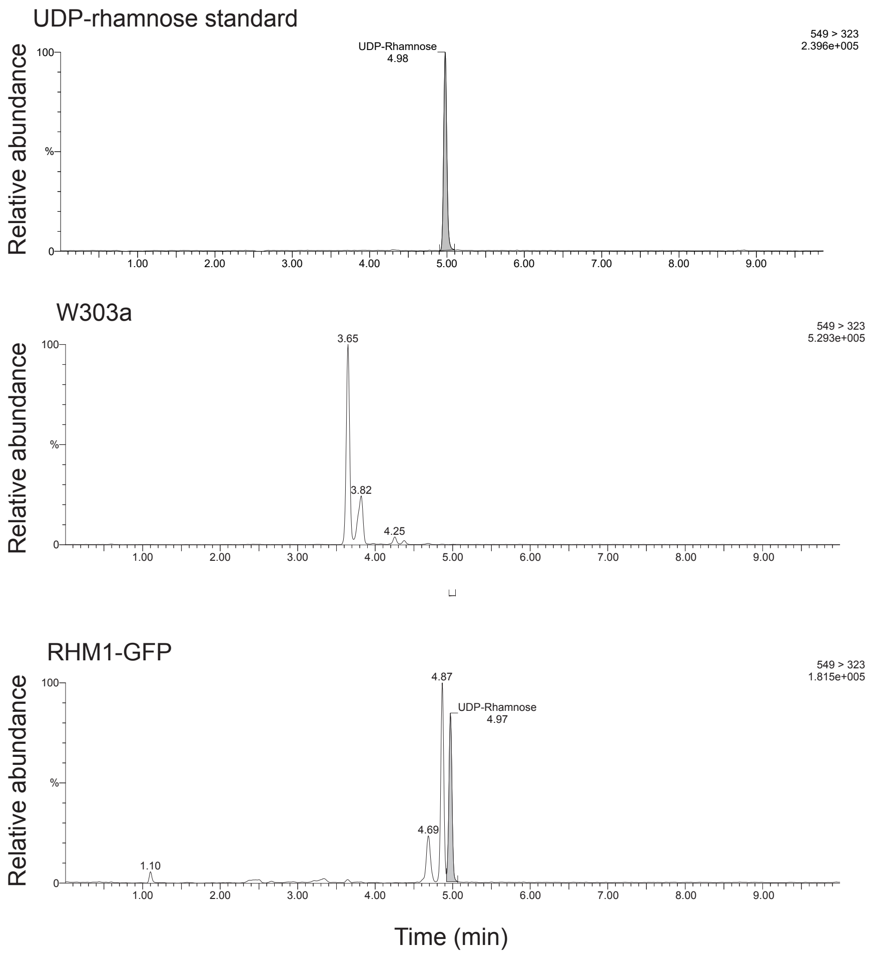


**Supplementary Figure 5. RHM1 is sufficient to synthesize UDP-rhamnose in yeast.** Liquid-chromatograph-mass spectrometry on a UDP-rhamnose standard (*top*), yeast lysate from W303a cells (*middle*), and lysate of W303a expressing RHM1-GFP (*bottom*). Grey peak denotes UDP-rhamnose. Molecular weight of UDP-rhamnose is 549.03 Da.

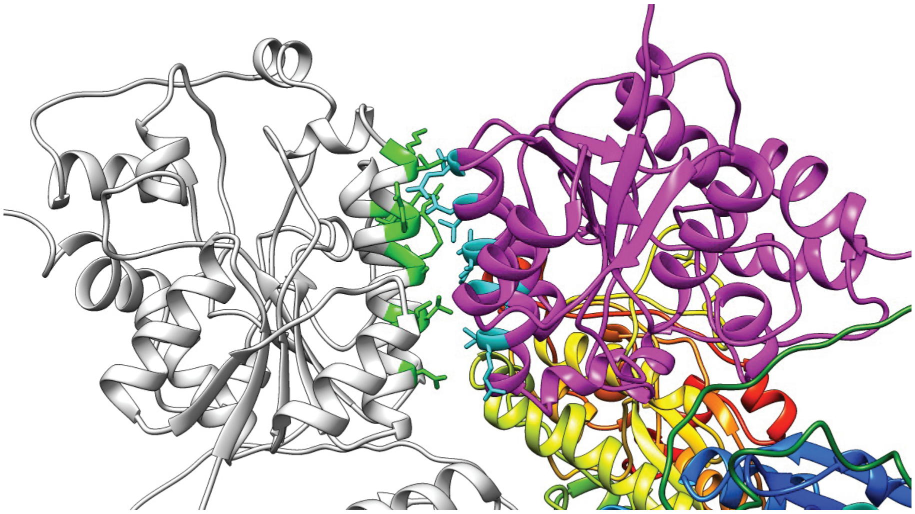

**Supplementary Figure 6. *In silico* prediction of RHM1-UER interaction.** Predicted interaction of UER1 (magenta) with a RHM1 homodimer (grey). UER1 is predicted to interact with BD2 of RHM1. Residues involved in the UER1-RHM1interaction are shown as cyan and green sticks.

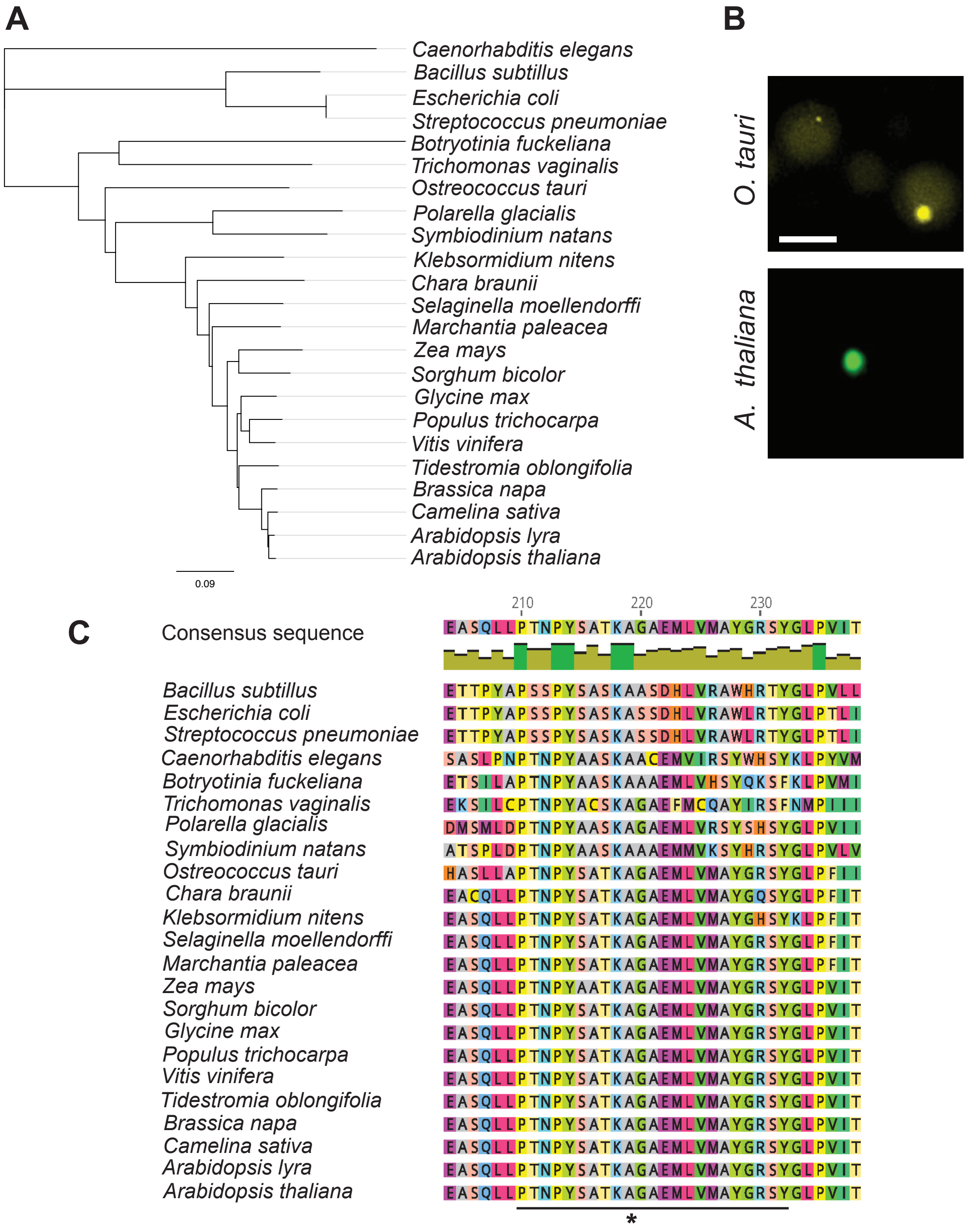


**Supplementary Figure 7. Evolutionary of RHM1 and rhamnosomes.**
**A**) Cladogram of the RHM1 amino acid sequence from rhamnose-synthesizing organisms, including examples from plants, invertebrates, protozoa, fungi, and bacteria. Scale bar denotes the number of substitutions per residue.
**B**) Micrograph of yeast expressing RHM1 from *Ostreococcus tauri* (*top*) and RHM1 from *Arabidopsis thaliana* (*bottom)*. Images are the same magnificaiton (scale bar = 5 µm). **C**) Protein sequence alignment of RHM1-like proteins from A, focusing on the domain implicated in RHM1 condensation. Asterisk marks A161 in *Arabidopsis thaliana*. Black line marks residues in the helix surrounding A161. Green bars below the consensus sequence indicate conserved residues.
